# Early Colonic and Microbial Responses Precede Hyperphagia in Short Bowel Syndrome: Insights from a Rat Model

**DOI:** 10.64898/2026.01.26.701700

**Authors:** Alice Garrigues, Mélanie Bourgin, Anne Dumay, Hounayda El-Jindi Shahrour, Maryline Roy, Alexandra Willemetz, Lara Ribeiro-Parenti, Nathalie Kapel, André Bado, Maude Le Gall, Johanne Le Beyec

**Affiliations:** UMR-S1149, Centre de recherche sur l’inflammation, INSERM, Université Paris Cite, Paris, France; AP-HP, Hôpital Bichat -Claude Bernard, Service de chirurgie Générale Oesogastrique et Bariatrique, Paris, France; AP-HP, Hôpital Beaujon, Service de Chirurgie colorectale et digestive, Service de Chirurgie digestive, oncologique et bariatrique, Clichy, France; AP-HP, Hôpital de la Pitié-Salpêtrière-Charles Foix, Service de Coprologie fonctionnelle, Paris, France; UMR-S 1139, INSERM, Université Paris Cite, Paris, France; Paris Center for Microbiome Medicine, Federation Hospitalo-Universitaire; Sorbonne Université, AP-HP, Hôpital de la Pitié-Salpêtrière-Charles Foix, Service de Biochimie Endocrinienne et Oncologique, Paris, France

**Keywords:** Short bowel syndrome, Intestinal adaptation, Gut microbiota, Hyperphagia, Colon in continuity, Enteroendocrine function

## Abstract

**Background:** Short bowel syndrome (SBS) resulting from extensive small bowel resection is characterized by severe malabsorption and represents the leading cause of intestinal failure. Although spontaneous intestinal adaptation can partially restore nutrient absorption, the temporal coordination and hierarchy of the adaptive mechanisms involved—particularly those linking the gut microbiota, enteroendocrine function, hyperphagia, and intestinal remodeling— remain incompletely understood.

**Methods:** We investigated the kinetics of spontaneous intestinal adaptation in a rat model mimicking type 2 SBS over a 28-day postoperative period. Body weight, food intake, gastrointestinal transit, fecal losses, intestinal morphology, enteroendocrine hormone secretion, hypothalamic neuropeptide expression, and gut microbiota composition were assessed longitudinally in SBS and SHAM-operated rats.

**Results:** Extensive small bowel resection induced marked early weight loss, accelerated intestinal transit, diarrhea, and increased fecal energy losses that persisted throughout the follow-up. Profound gut microbiota remodeling occurred as early as day 7, remained largely stable thereafter, and was characterized by reduced diversity and enrichment in Lactobacillaceae and Enterobacteriaceae. Early elongation of remaining colon and epithelial remodeling were observed, preceding the jejunal hyperplasia, which became evident from day 14 onward. Enteroendocrine adaptation was marked by an early increase in plasma peptide YY levels, whereas glucagon-like peptide-1 showed a modest response. Food intake was increased in SBS rats from day 7 onward, and hyperphagia developed gradually and reached a plateau by the end of the third postoperative week, in parallel with increased hypothalamic AgRP levels and reduced POMC levels. No significant improvement of intestinal transit and fecal energy losses was observed during the study period.

**Conclusion:** Intestinal adaptation to extensive resection follows a time-dependent sequence in which early gut microbiota remodeling and colonic adaptation precede hyperphagia and small intestinal remodeling. These findings highlight the gut microbiota and the colon as central components of the early post-resection adaptation and potential therapeutic targets in SBS.

## Introduction

Short Bowel Syndrome (SBS), the leading cause of chronic intestinal failure in adults ^1,2^ results from extensive resection of the small intestine, with or without concomitant resection of all or part of the colon. The remaining intestine no longer has sufficient capacity to ensure adequate nutrient absorption and maintenance of hydro-electrolyte balance, making long-term parenteral nutrition (PN) necessary. Although PN is life-saving, it is costly and invasive and, in the long term, is associated with significant morbidity and impaired quality of life ^3^

During the months to following intestinal resection, patients with SBS spontaneously develop a set of adaptive responses, which are largely stimulated by the resumption of oral or enteral nutrition and strongly influenced by the presence of the colon in digestive continuity. These adaptations progressively improve energy recovery and reduce dependence on PN, reflecting a gradual increase in the functional absorptive capacity of the remnant intestine ^1,4–6^. They are characterized by compensatory hyperphagia, which is considered beneficial because of its contribution to energy recovery, as well as by structural and functional modifications of the residual intestine and by profound remodeling of the gut microbiota ^1,2,6–8^. Increased efficiency of carbohydrate fermentation into short-chain fatty acids (SCFAs) in patients with SBS may further contribute to energy salvage ^9^.

However, these adaptations are heterogeneous and the underlying pathophysiological processes remain incompletely understood. Despite growing interest in intestinal adaptation, few studies have integrated the gut microbiota as an active contributor to post-resection absorption and functional recovery. We and others have demonstrated in individuals with SBS—including patients and rodent models—the development of colonic hyperplasia ^1,10–14^ and jejunal hyperplasia ^15^, thereby increasing the intestinal epithelial surface area available for nutrient exchange and absorption. This adaptive growth is associated with increased secretion of enteroendocrine peptides, particularly, those produced by distal L-type enteroendocrine cells. In particular, proglucagon-derived peptides (GLP-2 and GLP-1) and peptide YY (PYY) are known to promote intestinal trophicity (GLP-2), inhibit gastric emptying and slow intestinal transit. Profound alterations of the colonic microbiota are also observed in SBS patients and experimental models ^8,16,17^, characterized by an enrichment in *Lactobacillus* species—some of which are rarely detected in healthy adults ^8^—and depletion of obligate anaerobic microorganisms ^7,8,15,17^. We have shown that transplantation of this fecal microbiota, referred to as the *“lactobiota”*, from a SBS patient into germ-free rats induces colonic hyperplasia and increases GLP-1 and ghrelin secretion ^16^, suggesting a direct contribution of the microbiota to intestinal and hormonal adaptation. This *lactobiota* likely represents a reservoir of multiple and complex signals contributing to post-resection adaptive responses. Other groups have reported distinct—either beneficial or detrimental—effects on the health of SBS patients associated with the relative abundance of specific bacterial species within their microbiota ^18,19^, further supporting a link between intestinal adaptation and the microbiome in SBS.

Current knowledge of the pathophysiology of SBS and of intestinal adaptation over time relies largely on studies performed in animal models ^1,11,20^. Some studies have highlighted the contribution of the gut microbiota, hyperphagia, and enteroendocrine hormones to patients’ capacity for functional recovery. However, the temporal coordination of these adaptive components remains poorly understood. In particular, it is unclear whether microbiota remodeling, enteroendocrine responses, hyperphagia, and structural intestinal adaptation arise sequentially or concomitantly, and which of these processes may play a primary role in initiating or sustaining functional recovery. A clearer understanding of the temporal dynamics of these adaptive processes and of their potential interplay is therefore essential to optimize therapeutic strategies, reduce dependence on PN, and improve patient outcomes. The objective of this study was to investigate the kinetics of spontaneous intestinal adaptation following extensive intestinal resection, in order to define the temporal hierarchy of adaptive parameters and identify early events that may drive subsequent improvements in intestinal structure and function.

## Materials and Methods

### Approval and compliance

All experimental procedures were conducted in accordance with the European Community guidelines and were approved by the local ethics committee Paris Nord as well as by the French Ministry of Education and Research (APAFIS No. 38717).

### Animals

Male Wistar rats aged 7 to 11 weeks (n = 130), weighing between 250 and 400 g (Janvier Breeding Center, Le Genest-Saint-Isle, France), were used. Animals were housed in cages with ad libitum access to standard rat chow diet (C1314 Altromin, Genestil, Royaucourt, France) and water, and maintained under controlled temperature conditions (23 °C ± 1 °C) and a 12 h/12 h light–dark cycle (lights on from 7:00 a.m. to 7:00 p.m.). Animals were allowed to acclimate for 7 days prior to the start of the experiments.

### Surgical and experimental procedure

On the day of surgery, rats were randomly assigned to undergo either an 80% small intestinal resection including the ileum, ileocecal valve, and 20% of the colon followed by a jejuno-colonic anastomosis (SBS group), or a simple jejuno-jejunal transection with re-anastomosis (SHAM group) ^13,15^. Twelve hours prior to surgery, animals were fasted with ad libitum access to water. Baseline fecal and blood samples were collected the day before surgery (day 0). Animals were anesthetized by inhalation of 2.5% isoflurane (Iso Vet® 1000 mg/g). Standard aseptic procedures were followed throughout the surgery. In SBS rats, the intestine was resected from 25 cm distal to the ligament of Treitz to 3–4 cm beyond the cecum, followed by jejuno-colonic anastomosis. The lengths of resected and remaining intestinal segments (jejunum and colon) were measured in SBS rats. Jejunal and colonic segments from the resected intestine were collected for histological analyses, and mucosal scrapings were performed to collect epithelial cells for subsequent analyses.

To prevent postoperative pain and infection, animals received a subcutaneous injection of 1% Xylocaine (100 μL/100 g) (Astra, France) and penicillin G (20,000 units/kg) (Panpharma, Luitré, France). A subcutaneous injection of 10 mL of Bionolyte G5 (0.4% NaCl, 5.5% glucose, 0.2% KCl) (Fresenius Kabi, Louviers, France) was administered immediately postoperatively. Liquid diet (Nutrison, Nutricia, France) was available ad libitum 48 h after surgery, and ad libitum access to standard chow diet was possible from D4. Body weight and food intake were recorded daily for each rat (from day 1 and day 4 respectively).

Blood and fecal samples were collected weekly and stored at −80 °C for subsequent analyses. To investigate the kinetics of adaptive processes, rats were euthanized at 7-, 14-, 21-, or 28-days post-surgery. On the day before euthanasia, blood was collected from the tail vein into tubes containing a DPP-4 inhibitor (1:1,000; DPP-IV-010, Millipore, France). On the day of euthanasia, all animals were in the fed state with free access to water. Rats were anesthetized with isoflurane, and blood was collected by cardiac puncture into chilled tubes containing heparin and a DPP-IV inhibitor. Plasma was separated by centrifugation, aliquoted, and stored at −80 °C until analysis. Animals were then euthanized. Jejunal and colonic lengths were measured, and mucosal scrapings of the intestine and colon were performed for RNA analyses. The hypothalamus was also collected and rapidly frozen for total RNA extraction and RT-qPCR analyses. Intestinal and colonic segments were fixed overnight in paraformaldehyde (PFA) and stored in 75% ethanol at 4 °C until paraffin embedding for routine histological and immunohistochemical analyses. Cecal contents were collected at the level of the anastomosis for coprological and metabolomic analyses.

### Transit time study

Carmine red. Carmine powder (Sigma-Aldrich, C1022) was used as a gastrointestinal transit marker. A 6% carmine red solution was prepared by dissolving 6 g of powder in 100 mL of PBS containing 20% sucrose. A 100-μL aliquot of this solution was applied to a single food pellet, and all other pellets were removed from the feeder. The carmine-containing pellet was then provided to the animal. Once the pellet was consumed (typically within 10 min), the timer was started and stopped upon the appearance of the first red-colored feces. Transit time is reported in minutes.

### Determination of fecal energy losses and water content

Fecal samples stored at −80 °C were thawed at room temperature and homogenized. Samples from SHAM rats were diluted fivefold, whereas feces from SBS rats, which were liquid due to diarrhea, were analyzed undiluted. Nutrient composition (nitrogen, lipid, and carbohydrate content) was determined by near-infrared reflectance spectroscopy (NIR). Feces were then dried overnight in an oven to measure both wet and dry weights. The caloric content of the dried feces was determined using a bomb calorimeter.

### Measurement of plasma hormone concentrations

Plasma concentrations of PYY, GLP-1, and leptin were measured using MSD U-PLEX kits (Meso Scale Discovery, Maryland, USA). This technic allows customized multiplex immunoassays based on biotinylated capture antibodies coupled to specific plate spots. Target analytes bind to the capture antibodies, and electro-chemiluminescent detection antibodies complete the sandwich immunoassay. Signals generated upon application of an electrical potential were quantified using an MSD instrument, with signal intensity proportional to analyte concentration.

### Histological analyses

Sectioning was performed at the pathology platform of Bichat Hospital. Paraffin embedded formalin-fixed jejunum and colon segments were cut into 5-μm sections and stained with hematoxylin–eosin–saffron (HES). Slides were scanned, and villus height, crypt–villus height, crypt depth, muscularis thickness were quantified using QuPath software (version 0.4.1, open source). On average, 10 crypts (colon and jejunum) and 10 villi were analyzed per rat.

### Quantification of gene expression in jejunal and colonic mucosa and in the hypothalamus

Total RNA was extracted from 50–100 mg of jejunal or colonic mucosal cells using TRI Reagent (TRIzol, Invitrogen, Saint-Aubin, France) with a tungsten bead in a Tissue Lyser (30 Hz, 2 min). Chloroform (200 μL) was added, samples were centrifuged at 13,000 rpm for 15 min at 4 °C, and RNA was purified following the manufacturer’s instructions. The RNA pellet was resuspended in 60 μL RNase-free water and stored at −80 °C. RNA concentration and purity were measured using a NanoDrop spectrophotometer (NanoDrop™, Ozyme), and samples were diluted to 1 ng/μL for reverse transcription. cDNA was synthesized from 1 μg of RNA using the Verso cDNA Synthesis Kit (Thermo Fisher Scientific) according to the manufacturer’s protocol. cDNA was diluted 1:5 and then 1:100, and stored at −20 °C prior to quantitative PCR (qPCR). qPCR was performed in 96-well plates using the LightCycler® 480 system (Roche Diagnostics). The primers used are described in the supplemental data.

### Microbiota analyses

The intestinal microbiota was analyzed from fecal samples collected weekly over a one-month period. Samples were collected fresh, immediately frozen in liquid nitrogen, and stored at −80 °C until analysis. Total DNA was extracted from 200–300 mg fecal aliquots using the QIAamp Fast DNA Stool Mini Kit (Qiagen). DNA concentration and quality were assessed using a Qubit 4 fluorometer with the dsDNA HS Assay Kit (Life Technologies, USA). Microbiota composition and diversity were analyzed by high-throughput sequencing using an Illumina MiSeq platform. 16S rDNA amplicon reads were processed using the FROGS pipeline (Find Rapidly OTUs with Galaxy Solution) on the Galaxy platform, including read filtering, chimera removal, and clustering into operational taxonomic units (OTUs). Alpha diversity within groups was assessed using OTU richness (Observed) and the Shannon index, while beta diversity between groups was calculated using Jaccard and Bray–Curtis distances. Principal coordinates analysis (PCoA) was performed to visualize biodiversity distribution at the OTU level ^15^.

### Statistical analyses

Statistical analyses were performed using GraphPad Prism version 10 (GraphPad Software, San Diego, CA, USA). Values are expressed as mean ± SD or mean ± S.E.M, as indicated in the figure legends. Non-parametric tests were used: the Mann–Whitney test to compare two groups, the Kruskal–Wallis test followed by Dunn’s multiple comparisons test to compare more than two groups, paired, and two-way ANOVA with Bonferroni’s multiple comparisons test to compare multiple groups across different time points. A p value < 0.05 was considered statistically significant. Correlations were assessed using Spearman’s rank correlation test for non-parametric variables.

## Results

### Body weight evolution and assessment of motility and malabsorption

Daily body weight evolution is shown on figure 1A. As expected, SHAM rats lost some weight during the first two days following surgery (95.70 ± 0.30% of their preoperative weight, i.e., at day 0), then regained their weight and continued their normal growth until the 28th day (133.3 ± 2.12%). SBS rats lost an average of 15% (84.22 ± 0.42%) of their initial weight during the first four postoperative days, then their weight stabilized until day 7, and gradually increased between day 8 and day 28, reaching 96 ± 2.30% of their preoperative weight. At day 28, SBS rats had almost completely regained their initial body weight, but this recovery was heterogeneous (figure 1A). As a marker of the body weight loss in SBS rats we observed a decrease in plasma leptin concentration in SBS rats compared to SHAM at D7, D14 D21 and D28 (Supp. Figure 1).

**Figure 1:**
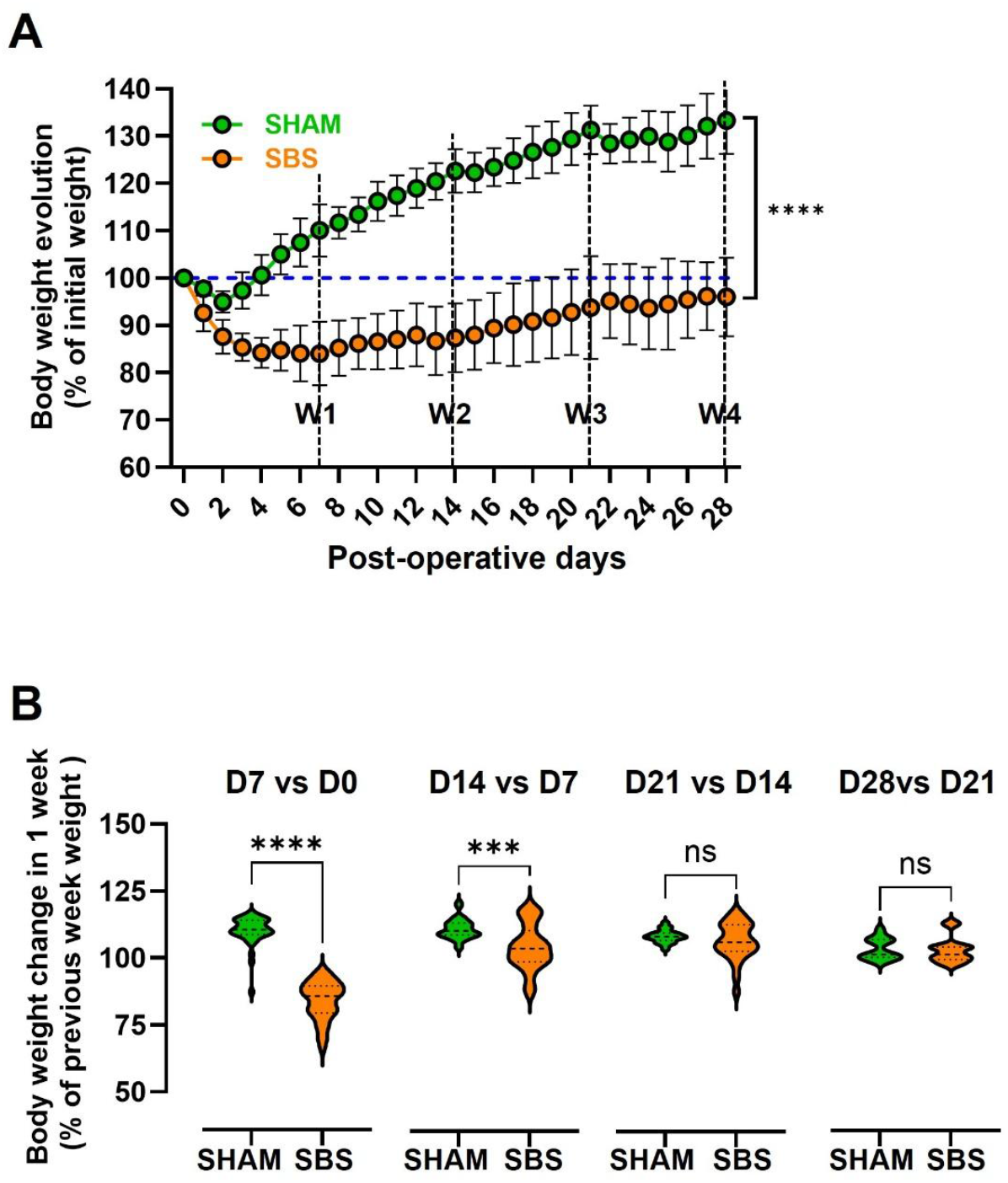
Time course of body weight changes following extensive small bowel resection. (A) Daily body weight evolution expressed as percentage of preoperative body weight (day 0) in SHAM (green) and SBS (orange) rats over the 28-day postoperative period (with n= 54-61 at D7, 40-44 at D14, 22-28 at D21, 12-15 at D28). Vertical dashed lines indicate weekly intervals (W1 to W4). Data are presented as mean ± SD. Statistical comparison used a Mixed-effects analysis. (B) Weekly body weight changes expressed as percentage of the body weight at the beginning of each week (D7 vs D0, D14 vs D7, D21 vs D14, and D28 vs D21) in SHAM (green) and SBS (Orange) rats. Individual values are shown with violin plots with n= 54-61 at D7, 40-44 at D14, 22-28 at D21, 12-15 at D28). ****p < 0.0001; ***p < 0.001 based on unpaired Mann Whitney test between groups for each time interval.

The curves of weight evolution (figure 1A) also suggested differences in the weekly weight change during week 1, 2, 3 and 4 in SBS rats. Comparing the percentage weight gain for each week of follow-up (weeks 1 to 4) in SBS and SHAM rats (Figure 1B), we observed significant differences in weekly weight variations during weeks 1 and 2, but as of the third week the weekly weight gain of SBS rats became similar to that of sham rats (+4.5% ±1.41 for SBS vs +8.8% ±2.38 for SHAM during the 3^rd^ week and +3.1% ±1.21 for SBS vs +3.6% ±1.14 for SHAM during the 4th week), suggesting that a new steady state had been reached in SBS rats.

Because SBS is associated with accelerated intestinal transit, which aggravates malabsorption and diarrhea, we investigated whether transit time improved over time in rats with SBS and we quantified the percentage of water in the feces (Supp. Figure 2A-C). The transit time of SHAM rats varied slightly over the 4 weeks (300 minutes on day 7 to 400 minutes on day 28) (Supp. figure 2A). Transit time in SBS rats was significantly reduced compared to SHAM rats but did not improve over time (Supp. Figure 2B). SHAM rats had very mild postoperative diarrhea in the first week, probably caused by the antibiotics used during surgery, and then had normal feces for the rest of the study (Supp. Figure 2C). SBS rats rapidly developed severe diarrhea, as indicated by a significantly increased water content in their feces, which remained high until day 28 (Supp. Figure 2D). We also evaluated the absorptive capacity of SBS and SHAM rats during this follow-up. Collecting feces over a 24-hour period, in order to obtain fecal output and loss over 24 hours relative to food intake, is complicated and unreliable in SBS rats. We therefore quantified fecal energy loss in lipids, nitrogen, and carbohydrates on fecal aliquots and expressed the results per gram of dry feces (Supp. Figure 2E-G). From day 7 onwards, SBS rats showed significantly higher lipid and nitrogen losses than SHAM rats (respectively p = 0.044 and p = 0.036 Mann-Whitney test SBS vs SHAM), with no improvement over time (p = 0.0364 on day 28 Mann-Whitney test SBS vs SHAM) (Supp. Figure 2E and 2F). Carbohydrate losses were not significantly higher in SBS rats, except on day 28 (p = 0.036, Mann-Whitney test SBS vs SHAM) (Supp. Figure 2G), consistent with what is observed in patients with SBS. We also analyzed mRNA levels of two key nutrient transporters, SGLT1 (sodium-glucose cotransporter) and PEPT1 (peptide transporter) in the jejunum (Supp. Figure 3A–D). SGLT1 mRNA expression was increased in SBS rats compared to SHAM rats at D14 and D21 (Supp. Figure 3A), and PEPT1 expression was increased at D21 (Supp. Figure 3C). In SBS rats, SGLT1 mRNA levels were significantly increased at D14 (p = 0.0306) and D21 (p = 0.0134) relative to D7, then returned to baseline at D28 (Supp. Figure 2B). PEPT1 mRNA levels were significantly increased at D21 (p = 0.0199 vs D7), then decreased again at D28 (Supp. Figure 3D). Increased expression of nutrient transporters such as SGLT1 and PEPT1 may have contributed to reducing sugar and protein losses in SBS rats; however, these differences observed at the mRNA level were modest.

### Changes in food intake and hypothalamic neuropeptides

Food intake was measured daily from Day 4 to Day 28 (figure 2A) since total free access to oral food (ad libitum) started on postoperative day 4. Food intake in SHAM rats remained stable (almost 30 g/day) throughout the monitoring period (figure 2A). In contrast, food intake in SBS rats increased as soon as ad libitum feeding resumed, exceeding that of SHAM rats from day 7 onward (33.6 g vs 27.6 p< 0.001 at day 7 based on Mann Whitney test). This food intake increased during 3 weeks and then stabilized reaching 52 ±3.87 g/day at day 28 which was significantly higher than that of SHAM rats (25.3 ±0.84 g/day) (p = 0.0004, Mann Whitney test), consistent with hyperphagia. At days 28, the cumulative food intake in SBS rats was significantly higher than in SHAM rats (p = 0.0012 vs SHAM, Mann Whitney test) (figure 2B). The weekly average food intake was not different between SBS and SHAM at the end of the first week, but it was during the other weeks (data not shown). Interestingly it increased significantly in SBS rats between the 1^st^ and the 2^nd^ week (p < 0.01) and between the 2^nd^ and 3^rd^ week (p < 0.05) but not between the 3^rd^ and 4^th^ week (p=0.65) (figure 2C), indicating that a plateau had been reached during the fourth week of follow-up.

**Figure 2.**
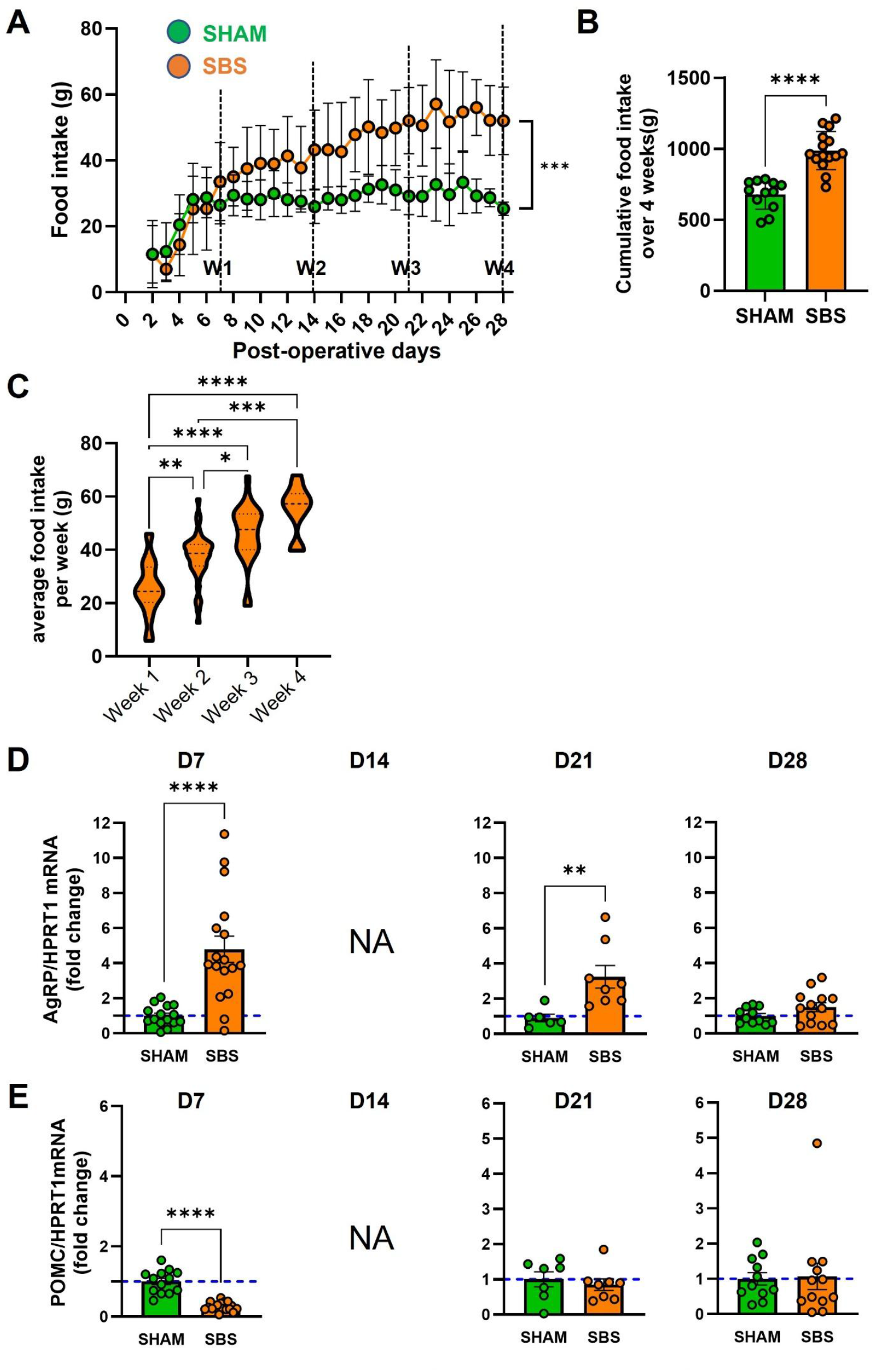
Development of hyperphagia and hypothalamic neuropeptide expression after small bowel resection. (A) Daily food intake measured from postoperative day 4 to day 28 in SHAM (green) and SBS (orange) rats until the 28th day (with n= 54-61 at D7, 40-44 at D14, 22-28 at D21, 12-15 at D28). Vertical dashed lines indicate weekly intervals (W1 to W4). Data are presented as mean ± SD. Statistical comparison used a Mixed-effects analysis. (B) Cumulative food intake over the 28-day follow-up period. Data are presented as mean ± SEM in a scatter plot. N= 12-15. Statistical comparisons were performed using unpaired Mann Whitney test. (C) Weekly average food intake in SBS rats during each postoperative week. Individual values are shown with violin plots with n = 42 at D7, 39 at D14, 28 at D21, 15 at D28). *p<0.05, **p<0.01, ***p<0.001, ****p<0.0001 based on unpaired Kruskal Wallis test followed by Dunn’s adjusted multiple comparison. (D, E) Hypothalamic mRNA expression in SBS (orange) compared to SHAM (green) rats at days 7, 21, and 28 after surgery of (D) the orexigenic neuropeptide AgRP and (E) the anorexigenic neuropeptide POMC, both normalized to hypothalamic HPRT1 mRNA level (delta CT). Data are presented as mean and individual values ± SEM. **p<0.01, ***p<0.001, ****p<0.0001 based on Mann Whitney test at each time point.

We then analyzed the mRNA levels of hypothalamic orexigenic (AgRP) and anorexigenic (POMC) neuropeptides at 1, 3, and 4 weeks of follow-up; measurements at week 2 were not available (figure 2D and 2E). At day 7, when food intake in SBS rats started to exceed that of SHAM rats, AgRP mRNA levels were 4.8-fold higher in SBS rats than in SHAM rats (p < 0.0001) (figure 2D) and POMC mRNA levels were 3.5-fold lower than in SHAM rats (p < 0.0001) (figure 2E). As a result, the AgRP/POMC ratio was markedly increased in SBS rats compared to SHAM rats (15.4-fold p < 0.0001), despite considerable inter-individual variability among SBS rats. At day 21, AgRP levels remained significantly higher in SBS rats than in SHAM rats (3.6-fold, p = 0.0023), whereas POMC mRNA levels no longer differed from between groups. Accordingly, the AgRP/POMC ratio was again significantly increased in SBS compared to SHAM rats (5.3-fold p = 0.0003). By day 28, there were no longer any differences in AgRP or POMC mRNA levels between SBS and SHAM rats. Overall, these hypothalamic changes are consistent with the temporal profile of food intake observed in SBS rats.

### intestinal adaptations

Intestinal adaptations, morphological evolution and endocrine functions, were assessed longitudinally over the postoperative period. The lengths of the small intestine and colon were measured at euthanasia at each time point (Figure 3). In SHAM rats the length of the small intestine decreased slightly over time, which was not the case in SBS rats (Figure 3A). Instead, SBS rats showed a tendency toward increased small intestinal length relative to the average length of intestine remaining just after resection (approximately 34 cm). In SBS rats the percentage of body weight regain relative to initial weight was positively correlated with small intestine length at days 21 and 28 (r = 0.683, p = 0.012 and r = 0.679, p = 0.025 respectively), whereas no such correlation was observed in SHAM rats. The length of the colon remained unchanged in SHAM rats throughout the follow-up period, whereas a significant colonic elongation was observed in SBS rats from day 14 onward compared with day 7, reaching a plateau by week 4. (Figure 3B).

**Figure 3.**
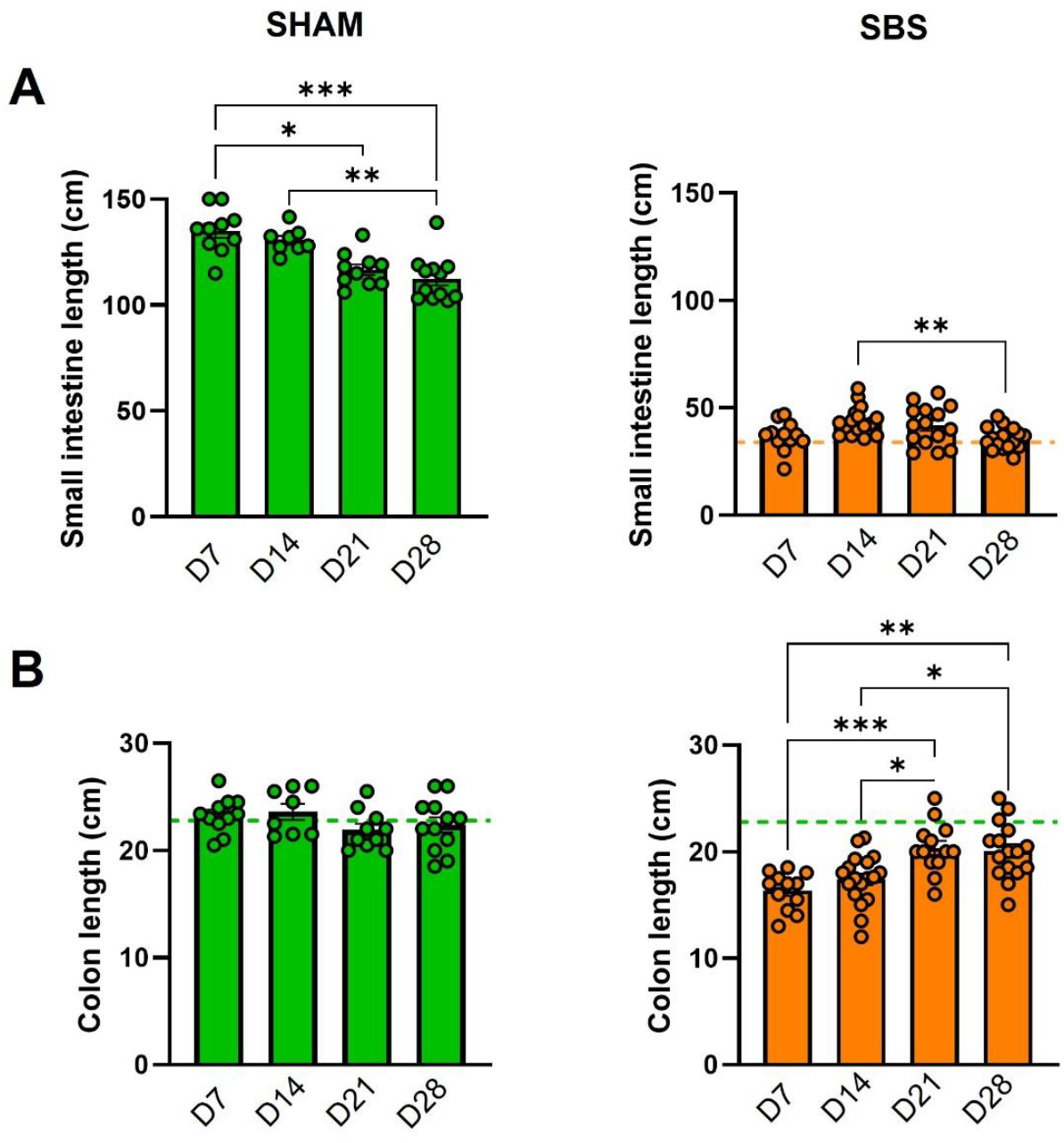
Adaptation of small intestinal and colonic length after extensive bowel resection. Comparison over time (days 7, 14, 21, and 28 post-surgery) of small intestine (A) and colon (B) Length evolution in cm within SHAM (green, n=8-12) or SBS (orange, n= 12-16) rats. Data are presented as mean ± SEM in a scatter plot. Data are presented as mean ± SEM in a scatter plot. N= 7 to 16, *p < 0.05; ** p < 0.01; *** p < 0.001 based on unpaired using a Kruskal Wallis test followed by Dunn’s adjusted multiple comparisons.

Morphological intestinal adaptation was further assessed by quantifying epithelial and muscular remodeling in the jejunum and colon (Figure 4). In the jejunum, the length of the crypto-villus axis increased in SBS rats from day 14 onwards compared with SHAM rats (p<0.01) (Figure 4A). This increase persisted until the end of the follow-up period, remaining approximately30-40% above SHAM rat values. In parallel, the thickness of the jejunal muscular layer was also significantly increased in SBS rats from day 14 onward (Figure 4B). In the colon, an increase in the depth of the colonic crypts (but not significant in this study) and the thickness of the muscularis (p<0.05) was observed from day 7 onwards (Figures 4C-D). The depth of the colonic crypts increased in SBS rats between D14 and D21 (p=0.03, Mann-Whitney) and remained elevated at D28 (Figure 4C).

**Figure 4.**
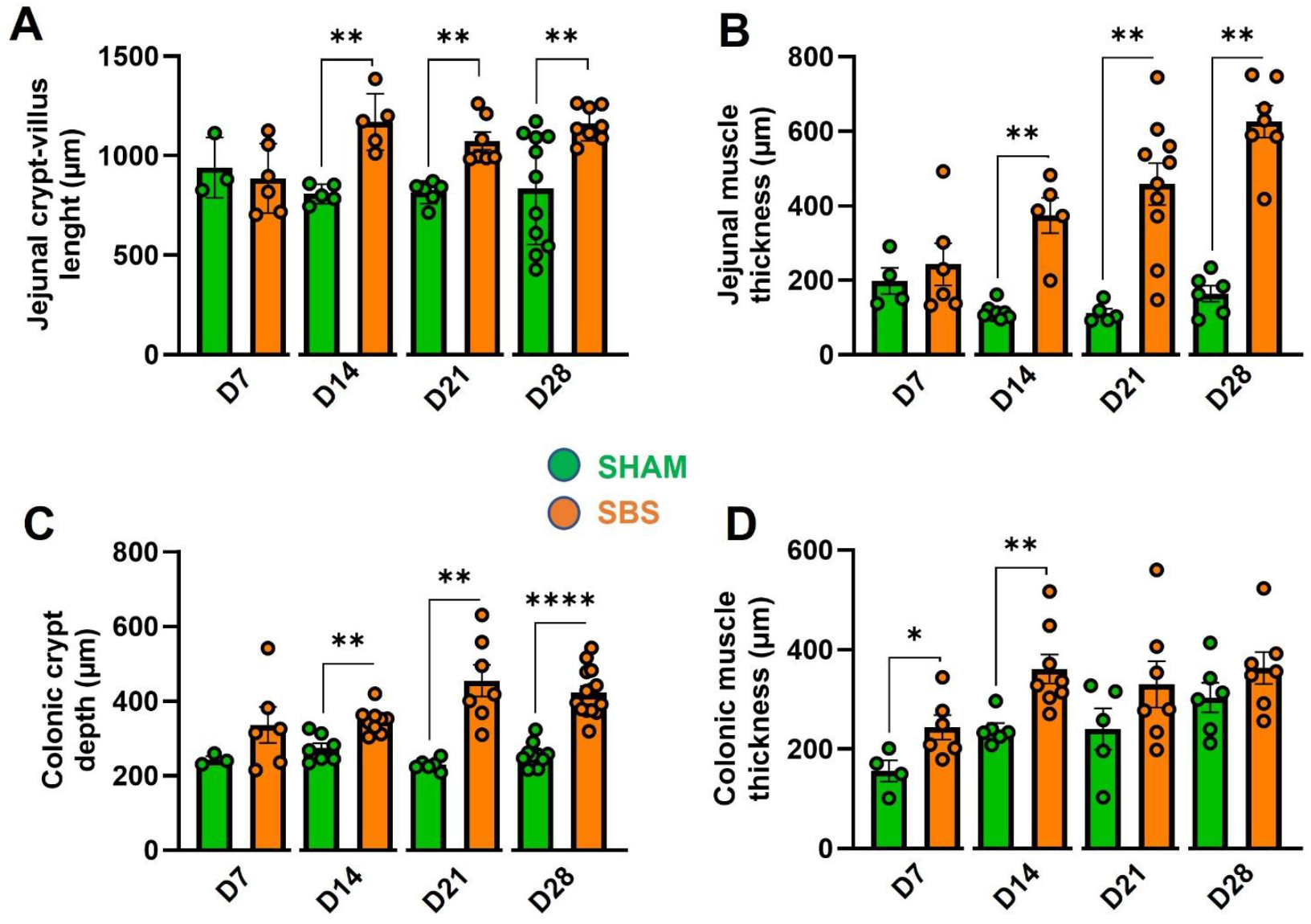
Structural remodeling of the intestinal wall following small bowel resection. Morphometric analyses were performed on hematoxylin–eosin–saffron–stained sections. Comparison in SHAM (n=3-10) and SBS (n=6-8) rats at each time of (A) length of the jejunal crypt–villus axis in μm, (B) thickness of the jejunal muscular layer in μm, (C) depth of colonic crypts in μm, D) thickness of the colonic muscularis. Data are presented as mean ± SEM, with individual values shown, * *p* < 0.05; ** *p* < 0.01; *** *p* < 0.001; **** *p* < 0.0001 vs. SHAM with based on unpaired Mann–Whitney test at each time point.

Therefore, SBS rats exhibited a rapid intestinal adaptive response, marked by early elongation of the colon and a tendency toward small intestine elongation, as well as an epithelial hyperplasia that seemed to begin first in the colon at day 7 and was clearly established in the jejunum and colon on day 14.

Intestinal endocrine function was assessed by analyzing the secretion and gene expression of hormones derived from L-type enteroendocrine cells (GLP-1 and PYY) (Figure 5). As previously shown, plasma PYY concentrations were significantly increased in SBS rats compared to SHAM rats from day 7 onwards, and remained increased at days 14 and 21, but not at day 28 (Figure 5A). Plasma GLP-1 concentrations did not differ significantly between groups, at day 7, although a trend toward increased levels was observed at days 14 and 21 (Figure 5B). Consistent with the plasma data, levels of colonic mRNA encoding PYY and proglucagon (precursor of GLP-1 and GLP-2) showed a tendency to increase from day 7 onward in SBS rats compared with SHAM rats. (Supp. Figure 4).

**Figure 5.**
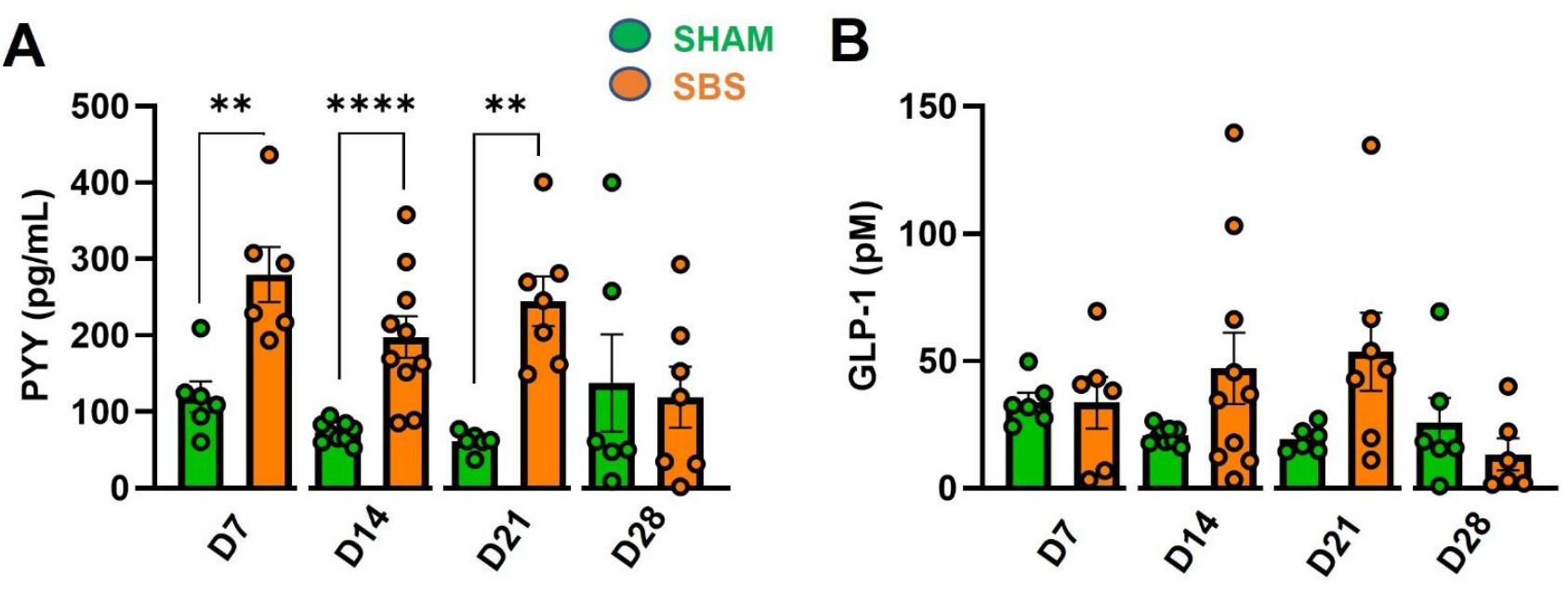
Changes in hormone secreted by L-enteroendocrine cell following small bowel resection. Comparison of plasma concentrations at each time point after surgery in SHAM (n= 6-9) and SBS (n= 6-10) rats of (A) peptide YY (PYY) in pg/ml and (B) glucagon-like peptide-1 (GLP-1) in pM, Blood samples were collected in the fed state. Data are presented as mean ± SEM, with individual values shown. * p < 0.05; ** p < 0.01; *** p < 0.001; **** p < 0.0001 SBS vs. SHAM based on unpaired Mann–Whitney test at each time point.

Together, these results indicate that enteroendocrine adaptation occurs early after extensive small bowel resection with a predominant and early PYY response, whereas GLP-1 changes appear more variable at the plasma level.

### Gut microbiota remodeling

The fecal microbiota was analyzed weekly from day 0 ‘before surgery), to day 28 (Figures 6). In SHAM rats, a decrease in alpha diversity was observed at day 7 (p = 0.0001), likely related to perioperative antibiotic treatment (Figure 6A). This decrease was transient, as alpha diversity returned to baseline from day 14 onward. Consistently, beta diversity was also altered at D7 and returned to baseline from D14 onward (Figure 6C). In SBS rats, alpha diversity was markedly reduced as early as day 7 (p = 0.0312) and remained severely reduced until day 28 (Figure 6B). Beta diversity was also significantly altered compared with the preoperative condition (D0) from day 7 onward (Figure 6D).

**Figure 6.**
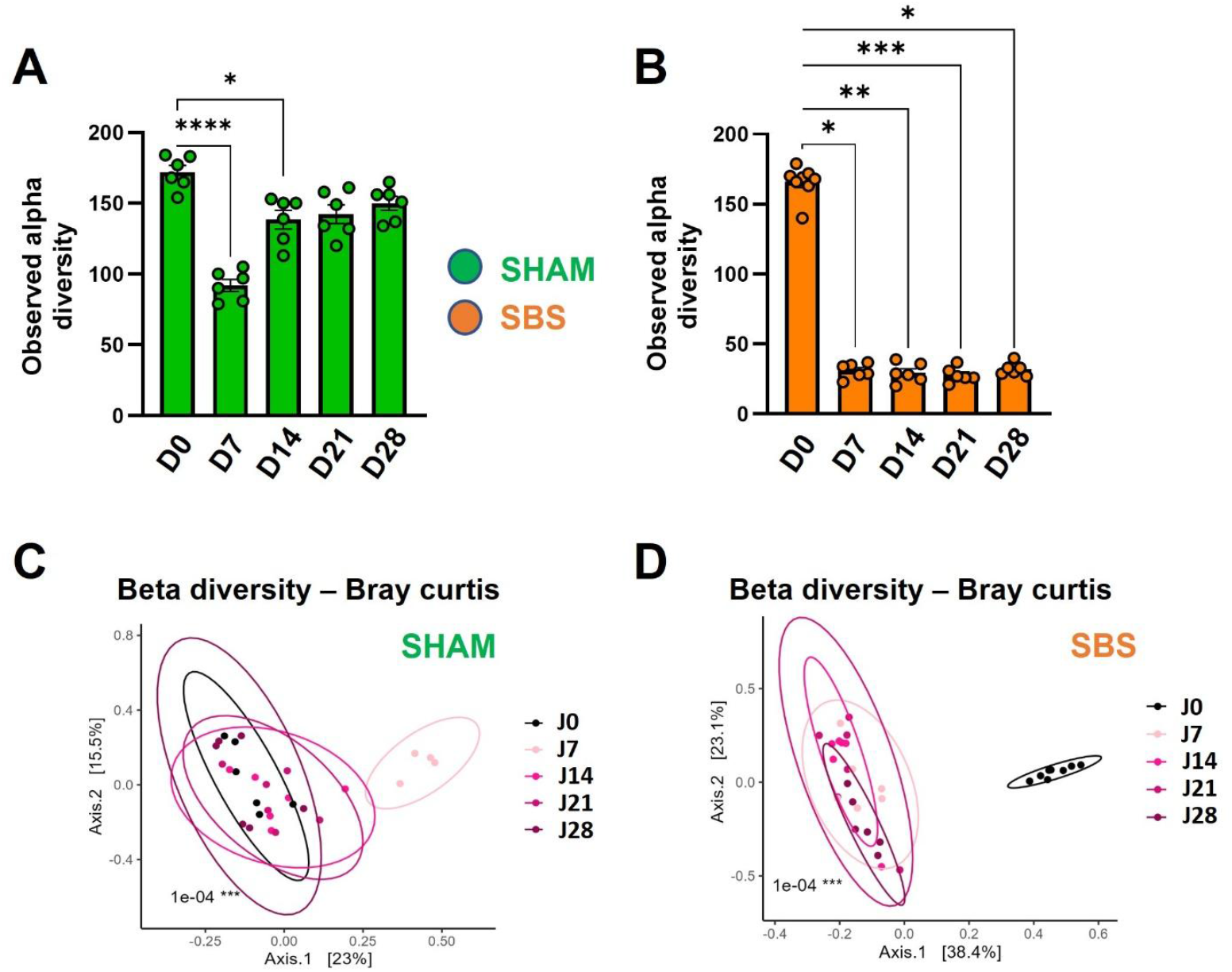
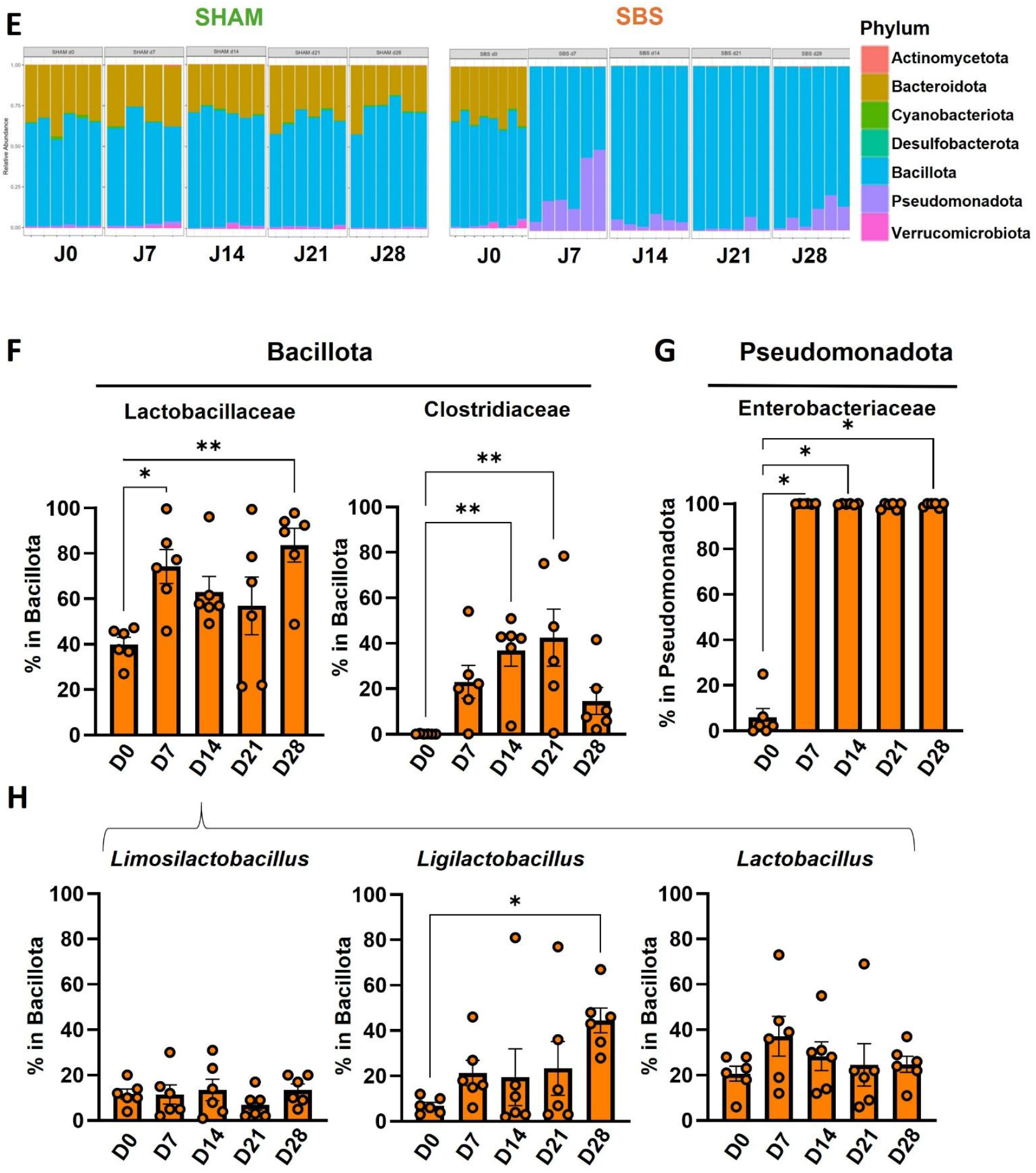
Time-dependent remodeling of the gut microbiota after extensive small bowel resection. (A–B) Alpha diversity of observed species in the gut microbiota estimated by observed OTUs counts in (A) SHAM (n=6) and (B) SBS (n=6) rats from day 0 (preoperative) to the 28th day post-surgery. (C–D) Beta diversity assessed by principal coordinates analysis (PCoA) based on Bray–Curtis distances in (C) SHAM (n=6) and (D) SBS (n=6) rats. P-value were determined using PERMANOVA test (Permutational Multivariate Analysis of Variance Using Distance Matrices). (E) Taxonomic composition and relative abundance of major bacterial phyla over time in SHAM (n=6) and SBS (n=6-8) rats and relative abundance of major bacterial families over time in SBS rats. (F) Relative abundance over time in SBS rats of two dominant bacterial families within the Bacillota phylum. (G) Relative abundance over time in SBS rats of Enterobacteriaceae within the Pseudomonadota phylum. (H) Relative abundance over time in SBS rats of *Lactobacillus, Ligilactobacillus*, and *Limosilactobacillus* genera within the Bacillota phylum. Data are presented as mean ± SEM, with individual values shown. p < 0.05; ** p < 0.01; *** p < 0.001; **** p < 0.0001 SBS vs. SHAM based in A, B, F, G and H on paired Friedman test with comparison to D0

The microbiota composition of SHAM rats did not exhibit major changes between day 0 and day 28. In contrast, SBS rats showed a marked decrease in Bacteroidota as early as day 7, associated with a sustained increase in Bacillota and Pseudomonadota throughout the study period (Figure 6E).

A detailed analysis of bacterial families within the Bacillota phylum (Figure 6F) revealed a significant increase in Lactobacillaceae proportion in SBS rats from day 7 (p = 0.03 vs. D0 by Wilcoxon test) and at D28 (p = 0.0013 vs. D0). The proportion of Clostridiaceae family also increased from day 7 (p = 0.06 vs. D0 by Wilcoxon test), but then decreased at D28. Enterobacteriaceae was the predominant family within the Pseudomonadota in the SBS rats (Figure 6G). Within the Lactobacillaceae family, *Lactobacillus* and *Ligilactobacillus* genera increased between day 0 and day 7 (respectively p = 0.09 and p = 0.06; Wilcoxon test), whereas *Limosilactobacillus* did not show a significant increase (Figure 6H). Other bacterial genera within the Bacillota, Bacteroidota were significantly decreased (data not shown).

## Discussion

In this study, we investigated the temporal dynamics of spontaneous intestinal adaptation following extensive small bowel resection in a rat model mimicking type 2 short bowel syndrome. By combining longitudinal analyses of body weight, food intake, intestinal morphology, enteroendocrine function, and gut microbiota composition, our work provides an integrated kinetic view of post-resection adaptation. While intestinal adaptation in SBS has been extensively described, most available data rely on cross-sectional analyses or focus on isolated parameters rather than their temporal coordination ^1,7,20^. Our results reveal that adaptation is a rapid but hierarchically organized process, in which early colonic (morphological and functional) and microbial responses precede the establishment of small intestinal structural remodeling and a central prioritization of energy intake is established to compensate for severe energy deficiency ^21^.

As expected, extensive small bowel resection induced marked early weight loss, accelerated intestinal transit, severe diarrhea, and increased fecal energy losses, reflecting an acute phase with profound malabsorption. Although SBS rats progressively regained body weight over the follow-up period, neither intestinal transit time nor fecal water content nor fecal energy losses showed clear improvement during the first month after surgery. These findings are consistent with previous observations in both experimental models and patients with SBS, in whom early adaptive responses do not immediately translate into normalization of digestive function ^6,12,20^. Modest increases in jejunal mRNA expression of nutrient transporters, including SGLT1 and PEPT1, were observed at intermediate time points. Similar upregulation of these transporters has been reported after massive bowel resection and is thought to partially compensate for the loss of absorptive surface ^22–24^. However, in our model, these changes remained limited and were not sufficient to normalize nutrient losses, suggesting that transporter regulation alone may play a secondary role during the early phase of adaptation. Moreover, as our analysis was restricted to mRNA expression, it does not necessarily reflect changes in protein abundance, membrane localization, or transporter activity, suggesting that regulation of nutrient transporters alone may play a secondary role during the early phase of adaptation.

A major adaptive response observed in SBS rats was the progressive development of marked hyperphagia. Food intake increased rapidly after the resumption of ad libitum feeding, with the most pronounced increase occurring during the second postoperative week, before stabilizing by the end of the follow-up period. Adaptive hyperphagia has been described as a key determinant of energy recovery in patients with SBS and in experimental models ^1,6,25^. In our study, the temporal profile of increased food intake closely paralleled changes in hypothalamic neuropeptide expression, characterized by increased AgRP and decreased POMC mRNA levels, as previously reported in models of jejuno-colonic SBS ^16,21^.

Morphological analyses revealed rapid but compartment-specific intestinal remodeling following resection. Elongation of the remaining colon and epithelial remodeling were detectable as early as day 7, whereas jejunal hyperplasia became clearly established from day 14 onward. These observations are consistent with previous studies reporting prominent colonic adaptation in SBS patients with colon in continuity and in rodent models ^10,12,26^. An increase in intestinal muscularis thickness was also observed, occurring early in the colon (day 7) and later in the jejunum (day 14). Intestinal muscular hypertrophy has received limited attention in studies of short bowel syndrome ^27^, and our observations may complement existing descriptions of morphological adaptation, suggesting that adaptive remodeling could extend beyond the mucosal compartment, including crypts and villi.

The early involvement of the colon suggests that it may serve as a key adaptive organ following extensive small bowel loss. Increased epithelial surface area, combined with enhanced microbial fermentation and enteroendocrine signaling, may contribute to improved energy salvage, particularly through short-chain fatty acid production ^9,26^. The positive correlation observed between small intestinal length and body weight regain at later time points further supports the functional relevance of structural adaptation, as previously reported in experimental SBS ^14,20^.

Enteroendocrine adaptation, particularly involving L-type enteroendocrine cells, was another early feature of the adaptive response. Plasma PYY concentrations were significantly increased from the first postoperative week onward, whereas GLP-1 levels showed a more modest and delayed increase. Elevated circulating levels of PYY and GLP-1 have been consistently reported in SBS patients with preserved colon and are thought to contribute to intestinal adaptation by modulating motility, secretion, and epithelial growth ^13,28,29^.

The early increase in PYY observed in our model may represent a compensatory “colonic brake” mechanism aimed at slowing gastrointestinal transit and optimizing nutrient contact time, even though no improvement in transit time was detected during the study period. In parallel, increased colonic expression of proglucagon and PYY mRNAs suggests enhanced enteroendocrine activity at the tissue level, in line with previous reports in both humans and rodents ^13,14^.

Hyperphagia occurred despite markedly elevated plasma PYY levels, which typically exert anorexigenic effects through Y2 receptors on arcuate AgRP/NPY neurons ^28,29^. This apparent discrepancy may reflect the predominance of the orexigenic central drive and/or a predominance of PYY (1–36), which lacks anorexigenic activity. Comparable elevations of PYY have been observed in SBS patients with colon in continuity, without consistent reduction in fecal output, highlighting variability in functional responses ^28^.

In both patients and experimental models of short bowel syndrome, circulating GLP-2 concentrations have been shown to increase early after resection, particularly when the colon is preserved ^28,30^. GLP-2 is a well-established intestinotrophic hormone that promotes mucosal growth and structural adaptation of the remnant intestine. Although GLP-2 was not directly assessed in the present study, activation of colonic L-cell pathways suggested by increased PYY and proglucagon expression may contribute to subsequent small intestinal adaptation.

One of the most striking findings of this kinetic study is the rapid and sustained remodeling of the gut microbiota following resection. SBS rats exhibited a dramatic reduction in microbial diversity as early as day 7, associated with a marked shift in microbial composition characterized by enrichment in Lactobacillaceae and Enterobacteriaceae and depletion of Bacteroidota. This microbial signature closely resembles that described in SBS patients and in other experimental models ^8,9,17^.

We have previously shown that transplantation of this SBS-associated microbiota, referred to as the “lactobiota,” into germ-free rats induces colonic hyperplasia and alters enteroendocrine hormone secretion such as proglucagon-derived peptides, supporting a direct role of the microbiota in post-resection adaptation ^16^. In this study we demonstrate that gut microbiota remodeling and early intestinal changes were largely established by the first postoperative week, whereas food intake, although already increased at day 7 in SBS rats, continued to intensify over the following weeks. This temporal dissociation suggests that increased food intake may not be the primary trigger of intestinal adaptation but contributes to amplifying and sustaining adaptive processes initiated ^6^. Another study revealed a positive correlation between food intake at postoperative day 15 and the relative abundance of Lactobacillaceae, two hallmark features of the SBS phenotype that displayed marked inter-individual variability. This observation is consistent with previous reports indicating that Lactobacillus strains can modulate eating behavior ^31,32^, further supporting a contributory role of this lactobiota in the integrated adaptive response following intestinal resection.

### An integrated kinetic model of adaptation in SBS

Taken together, our results support a model in which intestinal adaptation to extensive resection unfolds in a coordinated and time-dependent manner. Early after surgery, profound changes in the gut microbiota and the colon dominate the adaptive response, accompanied by increased secretion of enteroendocrine peptides. These early events likely create a permissive environment for subsequent structural remodeling of the small intestine and the development of hyperphagia, which together contribute to partial recovery of body weight and energy balance.

### Limitations and perspectives

This study has several limitations. Digestive efficiency was assessed indirectly, and longer follow-up periods would be required to determine whether the observed adaptive responses ultimately translate into improved nutrient absorption. In addition, although our kinetic approach highlights strong temporal associations between microbiota changes, hormonal responses, and structural adaptation, causal relationships cannot be definitively established.

Nevertheless, by integrating multiple adaptive parameters within the same longitudinal framework, this study provides new insights into the hierarchy of adaptive mechanisms in SBS. These findings reinforce the concept that the colon and the gut microbiota are early and central actors in post-resection adaptation and may represent promising therapeutic targets to enhance intestinal recovery and reduce long-term dependence on parenteral nutrition.

## Abbreviations

AgRP: Agouti-Related Peptide
DPPIV: Dipeptidyl Peptidase IV
GIP: Glucose-Dependent Insulinotropic Peptide
GLP-1: Glucagon-Like Peptide 1
GLP-2: Glucagon-Like Peptide 1
HPRT1: Hypoxanthine Phospho Ribosyl Transferase 1
PEPT1: Proton-Coupled Oligopeptide Transporter 1
POMC: Pro-Opiomelanocortin
SBS: Short Bowel Syndrome
SGLT1: Sodium-Glucose Cotransporter 1

## Author contributions

JLB, AB and MLG initiated and discussed the project. AG, MB, AB, MLG and JLB designed the experiments. AG, MB, AD, HS, MR, AW, LRP performed experiments and acquired data. AG, MB, AD, NK, AB, MLG and JLB analyzed and interpreted data. AG and JLB wrote the manuscript with inputs from AB.

All authors approved the final version.

## Acknowledgment

**We** are grateful to O. Thibaudeau and A. Chassac (Anapathology Platform Bichat Hospital) and Prof A. Couvelard (Anapathology Service Bichat Hospital) for help with IHC acquisition and interpretation. We also thank S. Fourati, C. Moulineuf, A. Goncalves, A. Mevel and S. Bouanane for their technical help and Prof. S Ledoux and Prof E. Larger for discussions and reviewing of the manuscript.

