## supplemental data for "Early Colonic and Microbial Responses Precede Hyperphagia in Short Bowel Syndrome: Insights from a Rat Model"

### Supplemental informations

Sequences of primers used

| Name | | Sequence | size |
|---|---|---|---|
| GCG | RatGcg_126_FWD<br>RatGcg_126_REV | TGAGATGAACACGATTCTCGAT<br>AAGATGGTTGTGAATGGTGAAA | 112 |
| SGLT1 | ratSGLT1_4_Left<br>ratSGLT1_4_Right | GAAGGGTGCATCGGAGAAG<br>CAATCAGCACGAGGATGAAC | 62 |
| PYY | ratPYY_17_Left<br>ratPYY_17_Right | TCCATCTCCTCCTGCTCATC<br>GAGCAGGACAAGCAGCATT | 100 |
| PEPT1 | ratPept1_20_Left<br>ratPept1_20_Right | AGGCATTTCCCAAGAGGAAC<br>CATTATCTTAATCTGCGAGATGAGC | 83 |
| AgRP | AgRP-FWD<br>AgRP-REV | CAGAGTTCTCAGGTCTAAGTC<br>TTGAAGAAGCGGCAGTAGCAC | 211 |
| POMC | ratPOMC_62_Left<br>ratPOMC_62_Right | AGGACCTCACCACGGAAAG<br>CCGAGAGGTCGAGTCTGC | 63 |

#### Supplemental Figures

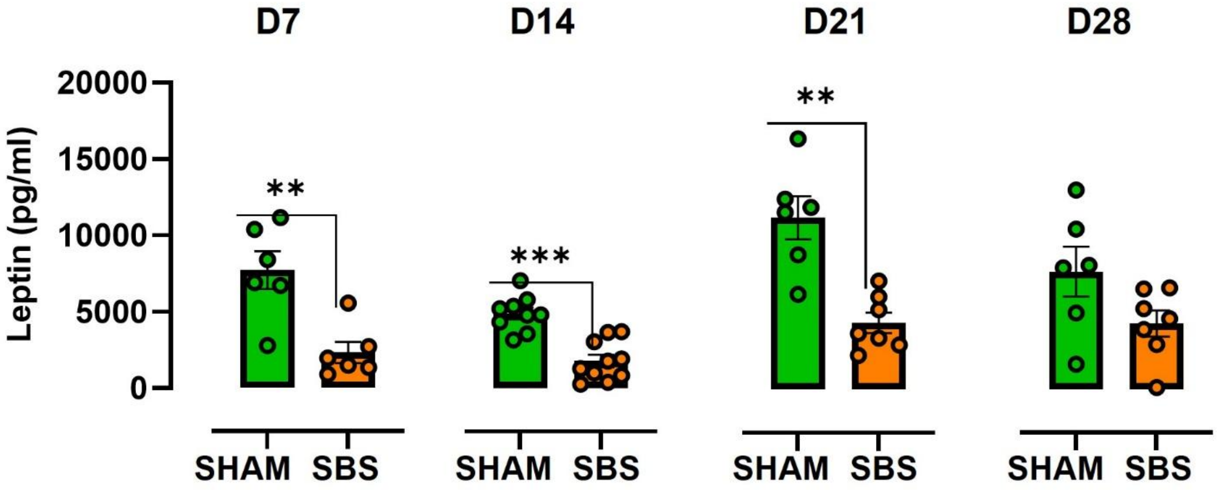


**Supplementary Figure 1. Changes in plasma leptin concentrations following extensive small bowel resection.** Plasma leptin concentrations in pg/ml at days 7, 14, 21, and 28 post-surgeries in SHAM (green) and SBS (orange) rats. Data are presented as mean ± SEM, with individual values shown. ** $p < 0.01$; *** $p < 0.001$ SBS vs. SHAM based on unpaired Mann–Whitney test at each time point

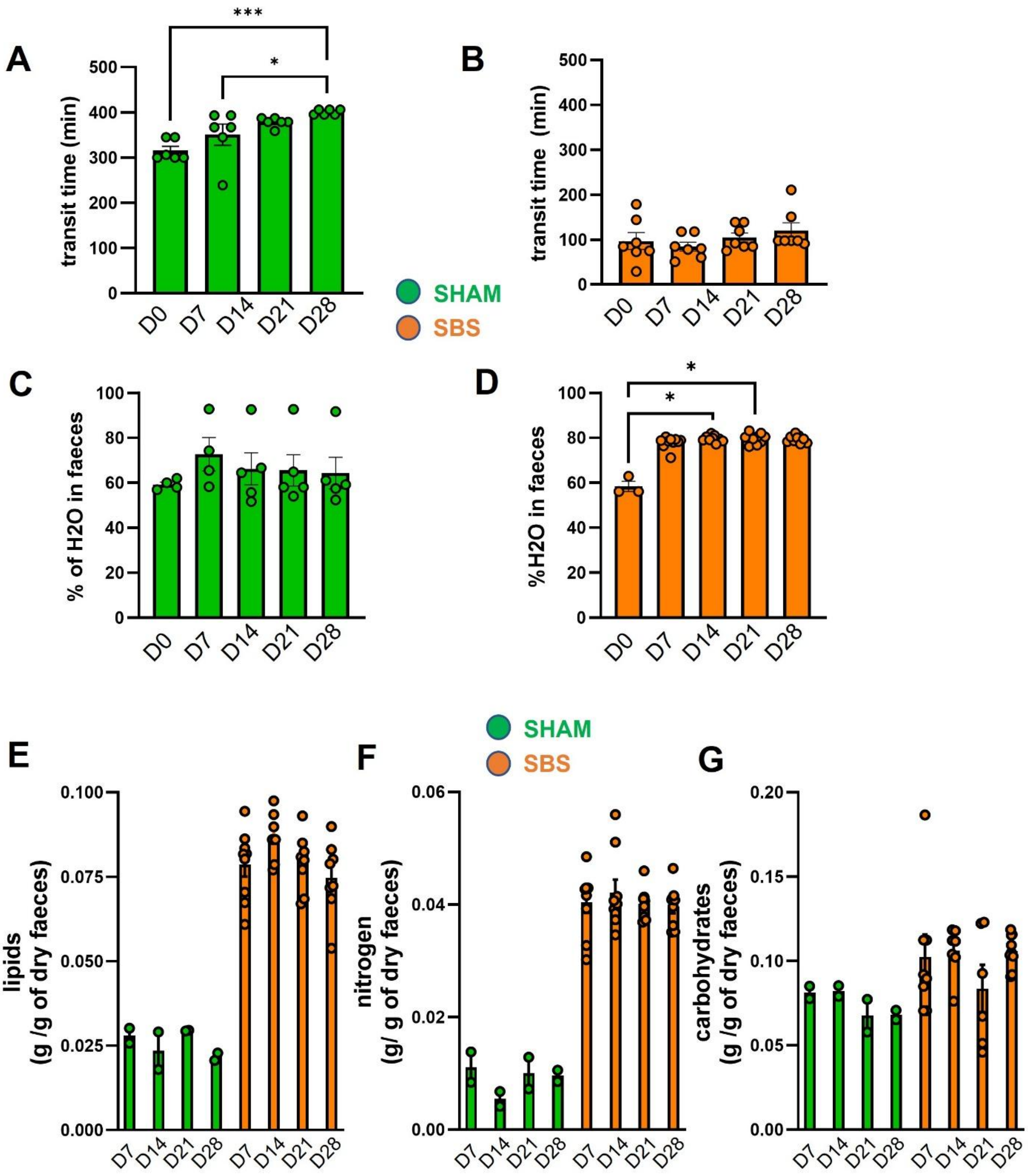


**Supplementary Figure 2. Persistent alterations of intestinal transit, fecal water content, and energy losses after small bowel resection.** (A, B) Gastrointestinal transit time in min measured at D0, D7, D14, D21 and D28 in (A) SHAM (green, n= 6) and (B) SBS (orange, n= 7) rats. (C-D) Fecal water content in percentage in (C) SHAM (green, n= 4-5) and (D) SBS (orange, n= 3-9) rats. Fecal losses in SHAM (green, n= 2) and SBS (orange, n= 9) rats expressed in gram per gram of dry feces. of (E) lipids, (F) nitrogen and (G) carbohydrates. Data are presented as mean ± SEM, with individual values shown. * $p<0.05$; *** $p < 0.001$ based on Kruskal Wallis test followed by Dunn’s adjusted multiple comparisons (A-D)

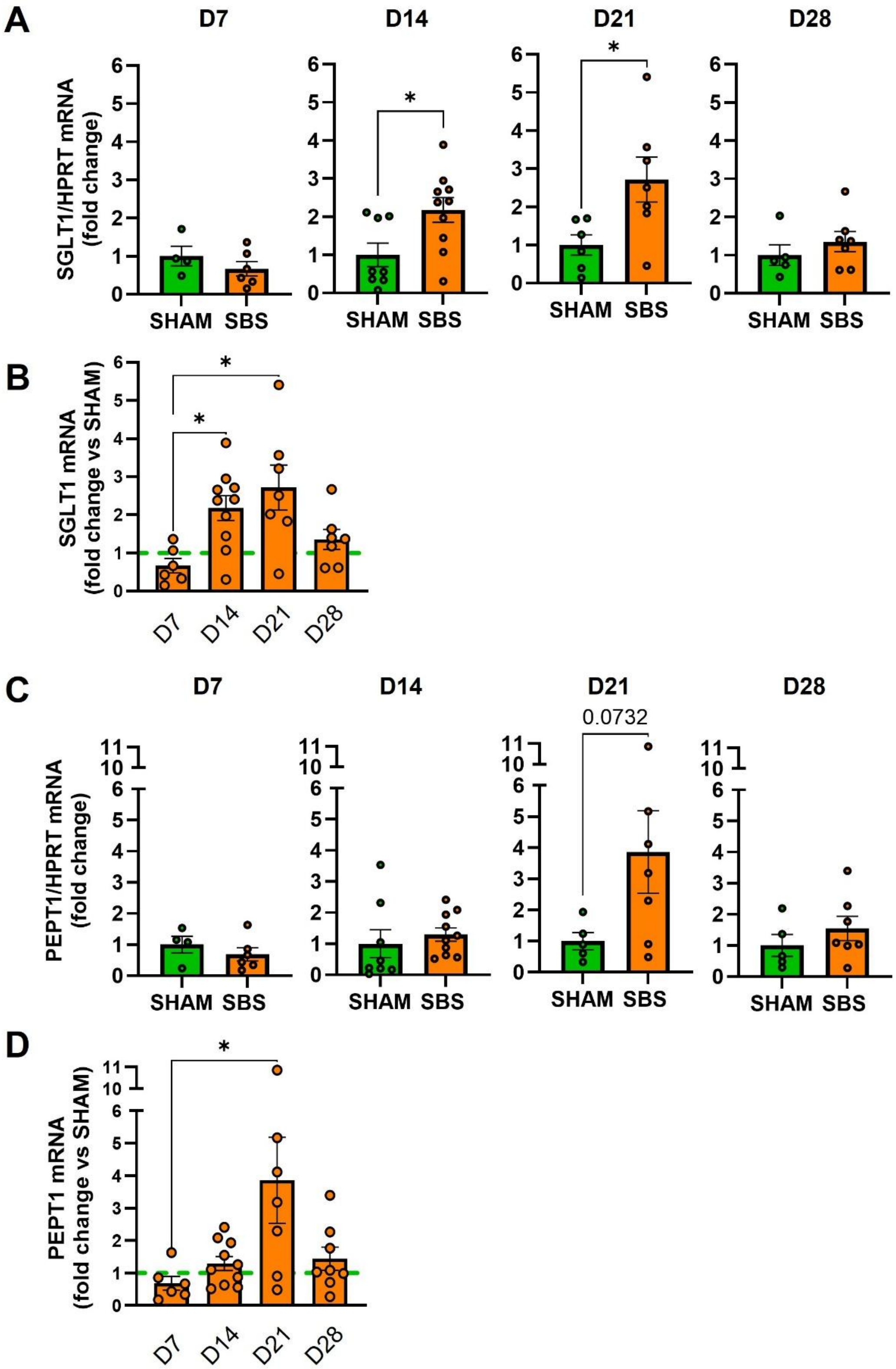


**Supplementary Figure 3. Time-dependent changes in jejunal nutrient transporter expression after small bowel resection.** (A–D) Jejunal mRNA expression at days 7, 14, 21, and 28 post-surgery in SHAM (green, n= 4-6) and SBS rats or within SBS (orange, n= 6-10) rats of (A-B) the sodium–glucose cotransporter SGLT1 and (C–D). Gene expression levels were normalized to housekeeping gene (L19) and expressed relative to the SHAM as the reference condition. Data are presented as mean ± SEM, with individual values shown. * $p < 0.05$ SBS vs. SHAM based on unpaired Mann–Whitney test at each time point (A and C) or in SBS rats over time based on Kruskal Wallis test followed by Dunn’s adjusted multiple comparisons (B and D)

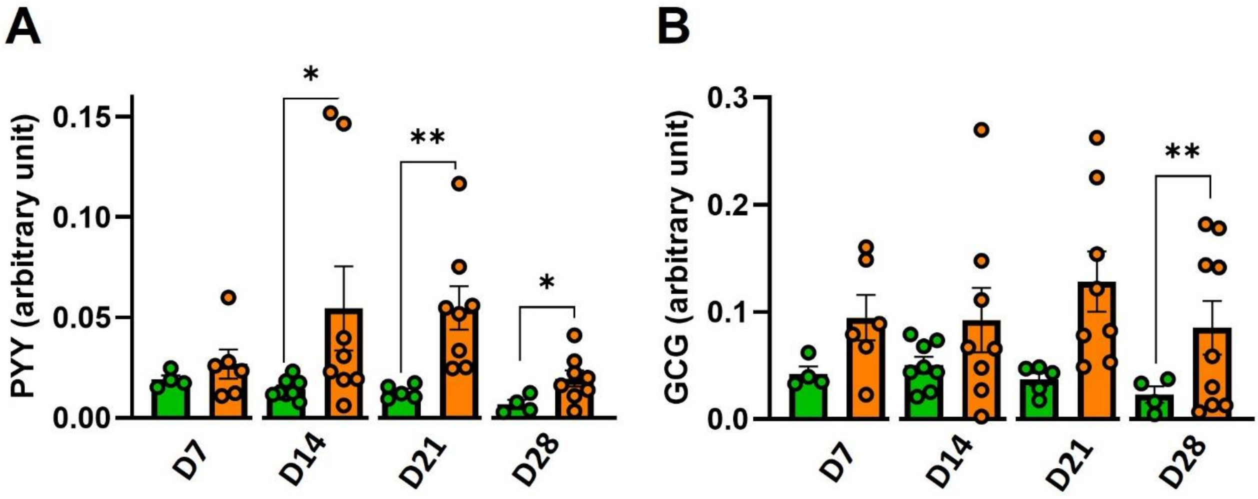


**Supplementary Figure 4. Colonic expression of enteroendocrine hormone genes following small bowel resection.** Colonic mRNA expression measured at days 7, 14, 21, and 28 post-surgeries in SHAM (Green; n= 4-8) and SBS (orange; n= 6-8) rats of (A) peptide YY (PYY) and (B) proglucagon (precursor of GLP-1 and GLP-2). Gene expression levels were normalized to housekeeping gene (L19). Data are presented as mean ± SEM, with individual values shown. * $p < 0.05$ ** $p < 0.01$ SBS vs. SHAM based on unpaired Mann–Whitney test at each time point.
